# Oxidative stress facilitates a diverse adaptive response in bacteria

**DOI:** 10.1101/2025.10.09.681464

**Authors:** Elise Bulaoro, Bharat Mishra, Michael J. McConnell, Anuradha Goswami

**Affiliations:** Department of Biological Sciences, University of Notre Dame, Notre Dame, US

## Abstract

The adaptive oxidative stress response allows bacteria to adapt to hostile environmental conditions. We explored how the evolution of cellular ROS in oxidative stress triggers physiological and genetic changes in multiple pathogenic species. Our study posits that growth-survival tradeoffs under oxidative stress enact a diverse array of genetic and physiological changes resulting in cross-resistance to acid, antimicrobial resistance, and metabolic reprogramming in bacteria.

## Oxidative stress impact on cell viability and antibiotic resistance

Alterations in the ecosystem or pristine environmental conditions could induce survival pressure in the microbial community ^1^. Central stress responses are often associated with the regulatory response in bacteria ^2^. Hence, it wouldn’t be surprising if the stress response to one predominant state (such as oxidative stress) could impose cross-resistance and co-selection. We hypothesize that when bacteria are subjected to oxidative stress conditions that result in loss of viability, global regulatory response pathways are activated. It enables bacteria to acquire enhanced stress tolerance, consequently contributing to the development of antimicrobial resistance (AMR). It is an ongoing debate whether exposure to antibiotics imposes oxidative stress in bacteria ^3-5^; however, to the best of our knowledge, no study has mechanistically integrated reactive oxygen species (ROS) causing oxidative stress and AMR in the absence of antibiotics in the environment.

Hydrogen peroxide (H_2_O_2_) is a naturally occurring stressor, produced as a by-product in bacterial metabolism, and dissociates to H_2_O_2_ → ^•^OH (ROS) + O_2_^6^. Consequently, bacteria have evolved to resist naturally occurring H_2_O_2_ stress levels ^7^. However, the increased use of H_2_O_2_ in the food industry, hospitals, wastewater treatment, and other applications can exert a selection pressure. Studying the effect of sub-MIC levels can be valuable to infer the effect of exposure to residual H_2_O_2_ on bacterial communities. The Fenton oxidation reaction (Fe^2+^ + H_2_O_2_ → Fe^3+^ + OH^-^ + ^•^OH) produces the highly reactive and non-selective hydroxyl radical ^•^OH (ROS) ^6^. Fenton oxidation is lethal to bacteria and was used in the study as a negative control for H_2_O_2_ oxidative stress. Comparing bacterial responses to these stressors can indicate whether regulatory responses are adaptable or endanger bacterial survival in a stressful environment.

*A. baumannii* ATC19606, *E. coli* MG1655, *P. aeruginosa* PAO1, and *S. aureus* USA300 represent diverse clinically and environmentally relevant isolates, and we therefore selected these isolates for a hypothesis-driven authors’ perspective. We subjected these strains to two different oxidative stress environments: a sub-minimal inhibitory concentration (sub-MIC) of H_2_O_2_ and sub-MIC H_2_O_2_ + Fe (III) (Fenton Oxidation Reaction) at a 1:1 molar ratio (H_2_O_2_: Fe (III)). Detailed information on research methodology is presented in the Supplemental file. Bacteria cultured in aerobic conditions were exposed to sub-lethal oxidative stress for 1 hour and 24 hours. The immediate phenotypic changes resulting from activated defense mechanisms were compared to control cultures (no stress). We observed that oxidative stress caused by either sole sub-MIC H_2_O_2_ or Fenton (sub-MIC H_2_O_2_ + Fe (III)) resulted in a substantial decline in the number of viable cells after a 1-hour exposure. The log fold changes in H_2_O_2_ and Fenton versus Control are given in Figure 1a [Average % reduction in colony forming unit, CFU/ml: *E*.*coli* MG1655 **H**_**2**_**O**_**2**_ = 53.4±6.2; **Fenton** = 31.9±26.6; *P. aeruginosa* PAO1 **H**_**2**_**O**_**2**_ = 59.8±27.13; **Fenton** = 99.8±0.05; *A*.*baumannii* ATC19606 **H**_**2**_**O**_**2**_ = 92.5±6.0; **Fenton** = 98.41±2.65; and *S. aureus* USA300 **H**_**2**_**O**_**2**_ = 94.11±6.50; **Fenton** = 99.88±0.18]. To evaluate the impact of oxidative stress on AMR, we performed a qualitative assessment of the antibiotic resistance profiles of the surviving bacterial colonies using the Kirby-Bauer disk diffusion method. Surviving colonies were tested against a panel of antibiotics, including chloramphenicol, erythromycin, kanamycin, rifampicin,tetracycline, and vancomycin.

**Figure 1.**
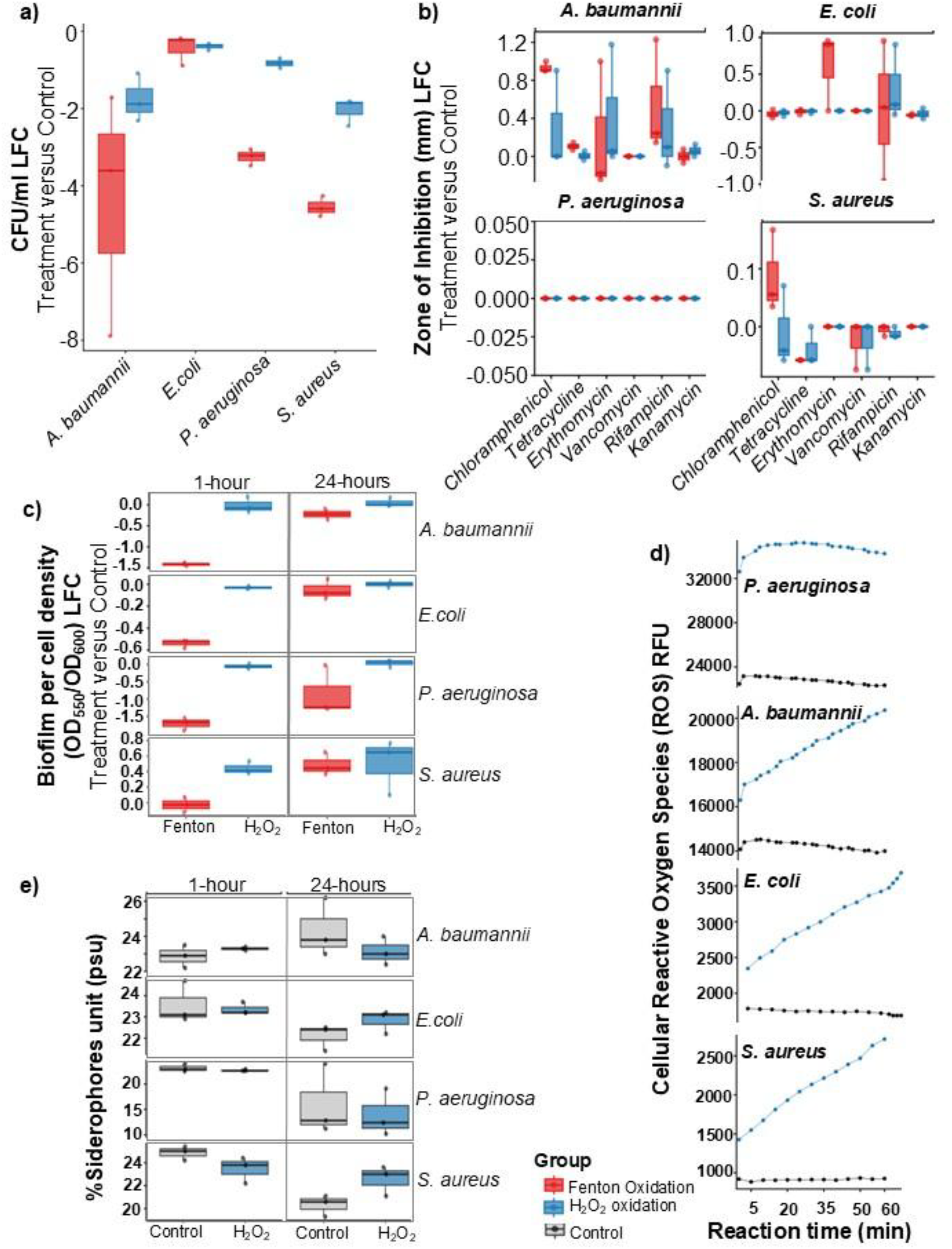
The Impact of Oxidative Stress on Bacterial Physiology - differential effects of two forms of oxidative stress—hydrogen peroxide (H_2_O_2_ oxidation and Fenton oxidation (H_2_O_2_ + FeCl_3_)—on four clinically relevant bacterial species: *Acinetobacter baumannii, Escherichia coli, Pseudomonas aeruginosa*, and *Staphylococcus aureus*. Red boxes and lines represent the Fenton oxidation treatment, while blue represents H_2_O_2_ oxidation. **a)** Change in Viability (CFU/mL LFC). Log Fold Change (LFC) in viable cell count (CFU/mL) of treated cultures compared to the untreated Control. Both treatments significantly reduce the viability of all strains, with the Fenton reaction showing a greater negative impact on *A. baumannii* and *S. aureus*. **b)** Change in Antibiotic Susceptibility (ZOI LFC). Log Fold Change in the zone of inhibition (ZOI) for treated cultures compared to untreated controls following exposure to six common antibiotics. For *A. baumannii* and *E. coli*, most treatments resulted in reduced ZOI (negative LFC), indicatingincreased resistance or reduced susceptibility. *P. aeruginosa* shows negligible change, while S. aureus displays mixed effects. **c)** Biofilm Formation (Log Fold Change). The LFC in normalized biofilm mass (OD550/OD600) after 1-hour and 24-hour exposure to treatments compared to the Control. *A. baumannii* and *E. coli* show reduced biofilm formation under both treatments at 24 hours. *P. aeruginosa* exhibits a sharp reduction in biofilm under Fenton oxidation, while *S. aureus* shows a significant increase in biofilm formation, particularly under H2O2 stress at 24 hours. **d)** Cellular Reactive Oxygen Species (ROS) Generation. Real-time quantification of intracellular ROS using a fluorescent probe over 60 minutes, expressed as Relative Fluorescence Units (RFU). All strains showed a sharp increase in ROS production upon exposure to the stress agents, confirming the induction of oxidative stress. *P. aeruginosa* and *A. baumannii* exhibit the highest overall RFU levels. **e)** Siderophore Production (% PSU). Quantification of siderophore production, expressed as Percent Siderophore Units (PSU), after 1-hour and 24-hour exposure to treatments compared to the Control. For most strains, the Fenton reaction (iron-limiting stress) induced a noticeable increase in siderophore production after 24 hours, particularly in A. baumannii and E. coli, consistent with a response to metal toxicity and iron deprivation.

Previous studies have suggested that, although oxidative stress can lead to cell death due to secondary killing, non-lethal levels of ROS may impact the development of antibiotic resistance in viable cells ^4^. Similarly, despite observing a reduction in overall bacterial populations following a 1-hour stress exposure, recovered colonies did not exhibit extreme shifts in phenotypic antibiotic resistance (measured with the Kirby-Bauer assay). It revealed a diverse relationship between stress and AMR in bacteria. While oxidation-treated bacteria mostly maintained their resistance, this suggests that existing defense mechanisms provide cross-protection. Albeit exceptional cases of increased susceptibility could occur, for example, *A. baumannii* ATCC 19606 and *E. coli* MG1655 became more susceptible to antibiotics, including erythromycin, tetracycline, rifampicin, and chloramphenicol, after exposure to stress. However, the sub-MIC levels used in this study (which are common in natural ecosystems) were not sufficient to cause a drastic, broad-spectrum shift in the existing resistance phenotypes of *P. aeruginosa* PA01 and *S. aureus* USA300, Figure 1 b. These data indicate that while the cell’s global survival strategy remains robust, the oxidative stress may have selectively compromised specific resistance pathways, highlighting a synergistic effect between stress and certain antibiotics. The ability of bacterial populations to recover and maintain their resistance phenotypes highlights the effectiveness of their diverse adaptive mechanisms, which allow them to endure prolonged oxidative stress conditions.

### Change in biofilm formation dynamics with oxidative stress exposure durations

The biofilm formation capacity, which is a major contributor to AMR, was assessed after 1 hour and 24 hours of exposure to H_2_O_2_ and Fenton, with relevance to acute or chronic responses. Overall, Fenton oxidation was observed to reduce biofilm formation after 1-hour of exposure; notably, an adaptive response was observed after prolonged exposure, where bacteria appeared to recover the biofilm loss in both oxidation treatments (Figure 1c). Both Fenton and H_2_O_2_ treatments led to a significant decrease in biofilm compared to the control as an acute response. However, the most notable response was observed in *S. aureus* USA300. While the Fenton treatment caused a strong decrease in biofilm after 1 hour, the H_2_O_2_ treatment led to a significant increase in biofilm formation at either 1 or 24 hours of exposure, highlighting a key difference in how these treatments interact with specific bacterial species over time.

### Phenotypic regulation in bacteria as a function of ROS damage

Estimating the rate of ROS produced inside bacteria can directly inform the associated regulatory response. The ROS kinetics of each strain were established by measuring cellular ROS (in Relative Fluorescence Unit (RFU)). We observed higher cellular ROS levels in bacteria when treated with H_2_O_2_ than in the controls (Figure 1d). All four strains followed zero-order kinetics of ROS formation irrespective of H_2_O_2_ exposure. These results highlight the biological bottleneck of ROS generation, suggesting that cellular ROS production is limited by the bacterial system, rather than controlled by H_2_O_2_ concentration. This contradicts the simple cause-and-effect relationship and opens an avenue to understanding the fundamental aspects of bacterial stress response.

Recently, a few studies have identified a bidirectional relationship between ROS and siderophores in bacteria ^8,9^. Fundamentally, siderophores are produced by bacteria in iron-limited environments; however, few studies have also indicated that they can also indirectly scavenge ROS by binding to surplus iron, thereby preventing iron-catalyzed oxidation and cell damage ^8^. To test the hypothesis, we quantified ROS production in cells exposed only to Fe and compared it with the control and H_2_O_2_-only treatment. The cellular ROS (RFU) produced by bacteria within 1 hour of incubation was significantly lower (T-test p-value = 0.03) in Fe alone treatment compared to H_2_O_2_ and control (no-spike). *E*.*coli* MG1655 (**H**_**2**_**O**_**2**_ = 29578; **Fe** = 3344; **Control=**17468), *P. aeruginosa* PAO1 (**H**_**2**_**O**_**2**_ = 34868; **Fe** = 38; **Control**= 22808), *A. baumannii* ATC1960 (**H**_**2**_**O**_**2**_ = 18783; **Fe** = 467; **Control**= 14266) and *S. aureus* USA300 (**H**_**2**_**O**_**2**_ = 2134; **Fe** = 414; **Control**= 913). These observations were in contradiction to classical Fenton chemistry, where iron acts as a catalyst to increase ROS. However, we anticipated that bacteria would regulate Fe through iron homeostasis by producing siderophores to avert excess Fe-intake and thereby reduce ROS.

Building upon our observation that siderophores could be involved in neutralizing ROS in a Fe-rich environment, we asked if siderophores were also produced in the absence of Fe when subjected to sub-MIC H_2_O_2_ oxidative stress. This will advance our understanding of bacterial adaptation mechanisms under oxidative stress conditions and the role of siderophores. Siderophore concentration quantified using a microtiter Chrome-Azole assay was increased in *S. aureus* USA300 after 1 hour exposure to H_2_O_2_ stress. A significant difference in siderophore formation after 24 hours of H_2_O_2_ treatment was measured; siderophores were increased in *P. aeruginosa* PAO1 and *A. baumannii* ATCC 19606 and decreased in *E. coli* MG1655 and *S. aureus* USA300 after prolonged stress (Figure 1e). Consistent with previous observations of biofilm formation, these findings reflect a diverse adaptive mechanism within the bacterial community, highlighting the need for further research to understand bacterial stress response and adaptation.

To unravel the regulatory response of bacteria under oxidative stress, we analyzed the publicly available RNA-Seq data of *E. coli* MG1655, which were exposed to 24 hours of H_2_O_2_-induced stress ^10^. Despite the genomic diversity in bacteria, *E. coli* MG1655 has a conserved oxidative regulatory response, making it an ideal model for this study^11^. The transcriptome profile demonstrated ∼ 75% variance among datasets, and some of the pathways enriched in H_2_O_2_-experiments highly expressed gene clusters related to anaerobic electron transport chain, fermentation, copper and silver ion transport and homeostasis, and acid fermentation (Figure, Figure S1a and b). We aimed to identify the key regulatory pathways in response to sustained exposure to H_2_O_2_ and shed light on the adaptive defensive mechanism against stress (details in Supplementary file). We found that prolonged H_2_O_2_ oxidative stress in *E. coli* MG1655 caused 4,396 genes to be differentially expressed, with 43 genes significantly upregulated and 14 genes significantly downregulated (*log2fc* ≥ |2|, *p*.*adj* < 0.1) (Figure 2a, Supplementary file). The transcriptomics data describes a multifaceted adaptive response in *E. coli* by reprogramming its cellular functions during exposure to oxidative stress (Figure S1b and Supplementary Table). The observed strategic shift involves the coordinated upregulation of protective mechanisms and the downregulation of resource-intensive processes (Figure 2a). The integration of prolonged H_2_O_2_ exposure transcriptome, gene regulatory, and protein-protein interaction networks in *E. coli* MG1655 resulted in a network with 426 nodes and 6,301 edges (connections including regulation and interaction) (Figure 2b; Supplementary Table).

**Figure 2.**
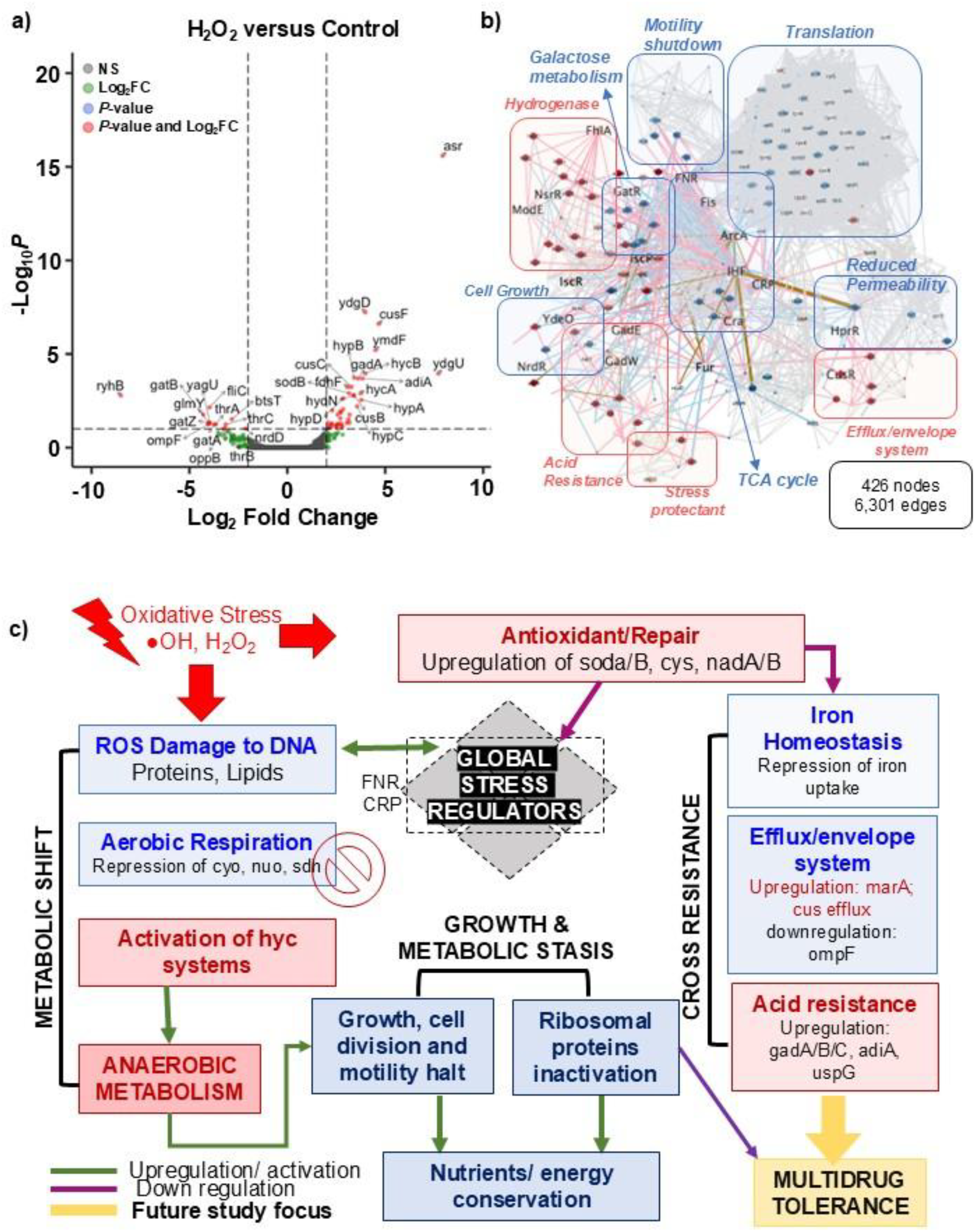
The regulatory programming of the H_2_0_2_ stressed *E. coli*. **a)** The volcano plot of differentially expressed genes (DEGs) for H202 vs Control *E. coli*. The significantly up- and down-regulated genes based on the threshold *log2fc* ≥ |2|, *p*.*adj* < 0.1 are represented in red. **b)** The integrated gene-regulatory network and protein-protein interaction (PPI) network (426 nodes, 6301 edges). The node type is circle = genes, rectangle = TF, triangle = regulator. The node colors are mapped with differential gene expression values (red = up regulation, blue = down regulation). The edge colors are mapped with type of priori interaction (+ = pink, -= blue, +-= brown, PPI = grey). The big node size are genes/TFs with high variance. The edge with is based on the edge-betweenness. The pathways are added on the network for easier network interpretation. **c)** The schematics representation of metabolic shift in *E. coli* after prolonged H_2_0_2_ exposure and their regulatory response.

### Regulatory network foundation to provide a framework for the potential functional paths and prospects of future study

#### Metabolic shift from aerobic to anaerobic

The most prominent adaptive strategy used by bacteria is a fundamental shift in metabolism. Global regulators such as FNR, ArcA, CRP, and Nac actively suppress genes involved in aerobic respiration and the tricarboxylic acid (TCA) cycle (cyo, nuo, and sdh operons). Downregulating these genes directly reduces the production of ROS, which are common byproducts of oxygen-dependent metabolism. Simultaneously, the cell activates anaerobic pathways. Regulators like FNR, ArcA, and YdeO upregulate genes for hydrogenases (hya and hyc systems). The extensive upregulation of genes involved in the formate hydrogenlyase (FHL) complex (fdhF, hycA-I, hydN, hypA-D) indicates a strong metabolic pivot. The shift towards less damaging, oxygen-independent pathways was observed as a critical defensive strategy.

#### Defense and Detoxification

Superoxide dismutases (sodA and sodB), widely known to neutralize harmful superoxide radicals ^12^ were upregulated with H_2_O_2_ exposure. The sodA and sodB expression are known to be repressed by both Fur (ferric uptake regulation) and the ArcA/ArcB (aerobic respiratory control) system, and activated by SoxR/SoxS (superoxide response) system^13^. In the regulatory gene network, we identified central role of Fur, FNR, Fis and ArcA genes, highlighting the cell’s sophisticated iron regulation and metabolic shifts as a consequence of oxidative response.

Antioxidant genes and chaperone proteins like cysA, slp, and uspG were upregulated. Activated glutamate decarboxylase and arginine decarboxylase systems (gadA-C, adiA), seen in this study, are known to provide acid resistance ^14^. Similarly, upregulation of nadA and nadB could ensure sufficient NAD^+^ in *E. coli* for metabolism and DNA repair, essential for offsetting DNA damage from oxidative stress. Most importantly, the upregulation of iraD, which stabilizes the master stress regulator RpoS, ensures a sustained and comprehensive adaptive response.

#### Resource Conservation and Growth Stasis

To survive prolonged stress, *E. coli* was observed to prioritize immediate survival over long-term growth by downregulating resource-intensive processes. The metabolic stasis by downregulation of cell division and amino-acid synthesis can render the antibiotics’ efficacy, as these are some of their most potent targets. Thereby, a coordinated adaptive response of bacteria against oxidative stress results in a lockdown strategy that can increase the risk of a broad-spectrum antibiotic predicament without the need for antibiotic encounter. The strategic trade-off was evident through reducing cell permeability, motility, and halting cell growth were predominantly observed in our study.

#### Reduced Permeability

Downregulation of outer membrane porins-OmpF, was regulated by regulators such as CpxR, OmpR, and Fur. It reduces cell permeability, creating a shield against external toxins and providing a form of antibiotic resistance. The downregulation of transporters like oppB also limits resource uptake. The Fis and CRP networks further reinforce this strategy by collectively downregulating ribosomal proteins (rpl and rps genes), halting protein synthesis, and conserving a significant cellular energy.

#### Cell growth arrest

The downregulation of genes for DNA synthesis (nrdD) and amino acid biosynthesis (thrA, B, C) indicates a pause in cell replication and growth. It could conserve energy and pivot resources to defense and repair. Reduced cell permeability, restricted transport, and DNA repair are shared mechanisms against antibiotics, which can result in multidrug resistance through oxidative stress-induced cross-resistance.

#### Motility Shutdown

The downregulation of genes for flagellar synthesis (fliC) and the transport of specific nutrients like pyruvate (btsT) and galactitol (gatA-Z) suggests a shift away from motility and specialized carbon sources.

## Conclusion

Our study reveals that sub-lethal H_2_O_2_ and potent Fenton oxidation drive bacterial evolution, while siderophores and iron could be facilitating the ROS defense. Transcriptomics and network systems biology approaches demonstrates that stress activates global regulators and shifts metabolism, leading to cross-resistance in *E. coli*. The paradigm shift in defense mechanisms suggests that future research should focus on dissecting the intricate connections between global stress responses and their commonality with multiple stress tolerance and AMR pathways.

## Author contributions

EB: Methodology, Data curation, Analysis, Writing – review; BM: Conceptualization, Methodology, Data curation, Analysis, Writing – review & editing; MM: Writing – review & editing; AG: Conceptualization, Methodology, Data curation, Analysis, Writing – Original draft, review & editing.

## References

1 Baquero, F. Environmental stress and evolvability in microbial systems. Clinical Microbiology and Infection 15, 5–10, doi:10.1111/j.1469-0691.2008.02677.x (2009).

2 Freyre-González, J. A., Alonso-Pavón, J. A., Treviño-Quintanilla, L. G. & Collado-Vides, J. Functional architecture of Escherichia coli: new insights provided by a natural decomposition approach. Genome Biology 9, R154, doi:10.1186/gb-2008-9-10-r154 (2008).

3 Dwyer, D. J. et al. Antibiotics induce redox-related physiological alterations as part of their lethality. Proceedings of the National Academy of Sciences 111, E2100–E2109, doi:10.1073/pnas.1401876111 (2014).

4 Qi, W., Jonker, M. J., Teichmann, L., Wortel, M. & ter Kuile, B. H. The influence of oxygen and oxidative stress on de novo acquisition of antibiotic resistance in E. coli and Lactobacillus lactis. BMC Microbiology 23, 279, doi:10.1186/s12866-023-03031-4 (2023).

5 Wong, F. et al. Reactive metabolic byproducts contribute to antibiotic lethality under anaerobic conditions. Molecular Cell 82, 3499–3512.e3410, doi:10.1016/j.molcel.2022.07.009 (2022).

6 Ofoedu, C. E. et al. Hydrogen Peroxide Effects on Natural-Sourced Polysacchrides: Free Radical Formation/Production, Degradation Process, and Reaction Mechanism—A Critical Synopsis. Foods 10 (2021).

7 Sen, A. & Imlay, J. A. How Microbes Defend Themselves From Incoming Hydrogen Peroxide. Front Immunol 12, 667343, doi:10.3389/fimmu.2021.667343 (2021).

8 Peralta, D. R. et al. Enterobactin as Part of the Oxidative Stress Response Repertoire. PLOS ONE 11, e0157799, doi:10.1371/journal.pone.0157799 (2016).

9 Arnold, E. Non-classical roles of bacterial siderophores in pathogenesis. Front Cell Infect Microbiol 14, 1465719, doi:10.3389/fcimb.2024.1465719 (2024).

10 Zorraquino, V., Kim, M., Rai, N. & Tagkopoulos, I. The Genetic and Transcriptional Basis of Short and Long Term Adaptation across Multiple Stresses in Escherichia coli. Mol Biol Evol 34, 707–717, doi:10.1093/molbev/msw269 (2017).

11 Choudhary, D., Foster, K. R. & Uphoff, S. The master regulator OxyR orchestrates bacterial oxidative stress response genes in space and time. Cell Systems 15, 1033–1045.e1036, doi:10.1016/j.cels.2024.10.003 (2024).

12 Carlioz, A. & Touati, D. Isolation of superoxide dismutase mutants in Escherichia coli: is superoxide dismutase necessary for aerobic life? The EMBO Journal 5, 623–630.

13 Fee, J. A. Regulation of sod genes in Escherichia coli: relevance to superoxide dismutase function. Mol Microbiol 5, 2599–2610, doi:10.1111/j.1365-2958.1991.tb01968.x (1991).

14 Richard, H. & Foster, J. W. Escherichia coli glutamate- and arginine-dependent acid resistance systems increase internal pH and reverse transmembrane potential. J Bacteriol 186, 6032–6041, doi:10.1128/JB.186.18.6032-6041.2004 (2004).

