## Supplementary for "Oxidative stress facilitates a diverse adaptive response in bacteria": Supplementary.pdf

**Supplementary File**

**Materials and Methods**

**Supplemental Figures**

### Materials and Methods

1. **Estimating the sub-Minimal Inhibitory concentration of H2O2 for the studied bacterial strains**

<u>Preparation of Dilutions and Inoculum</u>

Hydrogen peroxide ($H_2O_2$) working solutions were prepared by serial dilution in Mueller-Hinton Broth (MHB) or Luria-Bertani Broth (LB). An initial 2% (v/v) $H_2O_2$ stock was created by diluting 30% (v/v) $H_2O_2$. Two-fold serial dilution was then performed across 10 tubes by transferring 500 µL of the preceding concentration into 500 µL of sterile broth, resulting in a concentration range from 2.0% down to 0.00195% (v/v) in the dilution tubes. The molarity of the 1% (w/w) concentration, a mid-range dilution, was calculated to be approximately 0.294 M, assuming a solution density of 1.00 g/mL (Molar Mass $H_2O_2$ =34.014 g/mol).

Bacterial strains were grown in fresh MHB or LB for 2−4 hours at 37°C. The cultures were standardized spectrophotometrically to an OD600 between 0.08 and 0.12, corresponding to the 0.5 McFarland standard (approximately $1.5\times10^{8}$ CFU/mL). This standardized suspension was diluted 1:100 in fresh sterile broth to create the working inoculum at a final concentration of $1.5\times10^{6}$ CFU/mL.

<u>Microplate MIC Assay Setup and Incubation</u>

The $H_2O_2$ MIC assay was conducted in sterile 96-well microplates with three technical replicates per strain. Each well received 100 µL of the appropriate $H_2O_2$ dilution and 100 µL of the standardized bacterial working inoculum. The final H2O2 concentrations in the wells ranged from 1.0% (v/v) down to 0.00098% (v/v). Control wells included a Growth Control (GC) (100 µL MHB and 100 µL inoculum) and a Sterility Control (200 µL MHB only). After gentle mixing, the plates were sealed and incubated at 37°C for 18–24 hours.

MIC Determination

Bacterial growth was quantified by measuring the optical density at 600 nm (OD 600) using a microplate reader. An initial OD600 reading was taken immediately after plate setup (optional) and subtracted from the final OD600 reading to determine the net growth. The MIC was defined as the lowest concentration of $H_2O_2$ that resulted in an average Net OD value comparable to the Sterility Control wells and significantly reduced compared to the Growth Control, indicating the inhibition of visible bacterial growth.

2. **In Vitro Oxidative Stress Induction and Kinetic Analysis**

Bacterial Culture Preparation and Stress Agent Dosing

Bacterial cultures (e.g., E. coli MG1655, Acinetobacter baumannii ATCC 19606, Staphylococcus aureus USA300, and Pseudomonas aeruginosa PAO1) were initiated by diluting overnight stock cultures 1:100 (e.g., 150 µL into 14.85 mL) in sterile Luria-Bertani (LB) broth in 50 mL sterile tubes or flasks. All cultures were incubated at 37°C with shaking at 225 rpm for 3 hours to ensure cultures reached the mid-logarithmic growth phase before treatment initiation. For each bacterial strain, 15 mL cultures were divided into three treatment groups, each in technical triplicate (n=3), plus one media-only negative control:

i. Positive Control: Untreated culture.
ii. Treatment: Cultures exposed to H2O2 alone.

Fenton Treatment ($H_2O_2$ +$FeCl_3$): Cultures exposed to $H_2O_2$ and FeCl3 at a 1:1 molar ratio. The final concentrations of H2O2 used for each strain were determined based on preliminary MIC studies (e.g., E. coli: 0.029% $H_2O_2$; A. baumannii: 0.078% $H_2O_2$; S. aureus: 0.94% $H_2O_2$). For the Fenton treatments, $FeCl_3$ was added first from a 1 M stock, followed immediately by the required volume of 30% $H_2O_2$ to initiate the reaction. The final volume for all treated cultures was 10 mL.

Sampling and Growth Analysis

Kinetic sampling was performed at three time points: Baseline (T=0 h): Immediately before the addition of $H_2O_2$ and $FeCl_3$, a 5 mL aliquot was withdrawn from each culture for baseline analysis. Short-term Exposure (T=1 h): After 1 hour of incubation post-treatment, a 5 mL aliquot was withdrawn from all treatment and control tubes. Overnight Exposure (T=24 h): The remaining cultures were incubated for a total of 24 hours, after which a final 5 mL aliquot was withdrawn. Bacterial growth kinetics at all time points were monitored by measuring the optical density (OD) in a 96-well microplate format (OD 600) using a plate reader. The collected 5 mL aliquots were also immediately processed for downstream analyses (e.g., cell viability, ROS measurement, etc.).

3. **Viable Cell Counts (CFU/ml)**

Bacterial cell viability was determined by the standard plating method[1]. The samples were serially diluted and 10µl aliquots of selected dilutions were spread onto the surface of appropriate agar plates and incubated at the optimal growth temperature. Colonies forming on plates (Colony Forming Units/ ml) were enumerated. The cell concentration was calculated using: CFU/ml = Number of colonies × dilution factor/ Volume plated [1].

4. **Antimicrobial Susceptibility Testing (Kirby-Bauer)**

Antimicrobial susceptibility was assessed using the Kirby-Bauer disk diffusion method according to standard protocol [2]. The 10µL of bacterial culture in broth was added to Mueller-Hinton (MH) agar plates and a sterile cotton swab was used to create a uniform lawn of bacteria over the entire surface. Sterile antibiotic-impregnated disks were placed onto the inoculated agar. Plates were inverted and incubated at 37°C for 24 hours. Susceptibility was determined by measuring the diameter of the zone of inhibition (ZOI) around each disk and comparing these values to interpretative standards.

5. **Biofilm Quantification (Crystal Violet Assay)**

Biofilm biomass was quantified using the Crystal Violet staining method in -well microtiter plates[3]. The washed pellet after treatment was processed for biofilm assay. Pellets were incubated in LB broth in a 96-well plate to grow biofilm. After a 24-hour incubation period for biofilm formation, planktonic cells were gently removed, and the adherent biofilm was fixed using methanol or ethanol. The plates were then stained with 0.1% crystal violet solution and incubated at room temperature for 20 min. Unbound dye was washed off using distilled water, and the plates were allowed to air-dry. The bound dye (proportional to total biofilm biomass) was solubilized using 30% acetic acid, and the absorbance of the resulting solution was measured using a microplate reader at OD 550nm.

6. **Intracellular Reactive Oxygen Species (ROS) Measurement**

Intracellular levels were measured using the Invitrogen Thermo Fisher Sientific Reactive Oxygen Species (ROS) Fluorometric Assay Kit (Green). Bacterial cells were incubated with fluorescence dye (2',7'-dichlorodihydrofluorescein diacetate (DCFH-DA)) and grown to reach log-phase to incorporate dye into the bacterial cells. The extra dye in the media was then washed, and cells with dye were incubated in a medium spiked with sub-MIC $H_2O_2$.

Bacterial cells were incubated in the dark as described in kit protocol. After washing, fluorescence was measured using a microplate reader with an excitation wavelength at 488nm and an emission wavelength at 525 nm.

7. **Siderophore Detection (Chrome Azurol S - CAS Method)**

Siderophore production was detected using the universal colorimetric Chrome Azurol S (CAS) assay [4]. The reagent consists of a blue complex formed between the dye, hexadecyltrimethylammonium bromide, and ferric iron. Bacterial culture supernatants collected before washing after the treatment were mixed with the CAS solution. If siderophores are present, they scavenge the iron from the complex, resulting in a color change from blue to orange/yellow. Siderophore activity was quantified by measuring the absorbance of the supernatant-CAS mixture at 630 nm.

Siderophore production is inversely proportional to the absorbance at 630 nm. Higher siderophore concentration indicates more iron removal from the CAS, leading to a decrease in blue color and thus lower absorbance.

*The percentage of siderophore units can be calculated using the formula:*

***Siderophore Unit (%) =*** $[(A_{control} - A_{sample})/A_{control}] \times 100$

*Where* $A_{control}$ *is the absorbance of the negative control (CAS + LB) and* $A_{sample}$ *is the absorbance of the sample (CAS + supernatant).*

**Bulk transcriptome analysis**

The raw RNA-Seq data of *E. coli* multi stressor experiment was downloaded from NCBI GEO id GSE58325 [5]. Secondary analysis of bulk RNA-seq FASTQ files was performed with the nf-core/rnaseq pipeline[6] with the latest reference genome. The analysis includes initial QC, trimming, alignment, quantification and post-alignment QC. Tertiary analysis including QC, normalization, and differential gene expression analysis between pairwise comparisons was performed by the DESeq2 [7]. Standard cutoff i.e. *log2fc* ≥ |2| and *adjusted P-value* by FDR < 0.1 was selected for DEGs quantification. The *k-mean* clustering was performed by iDEP[5]. The gene ontology and enrichment analysis was performed by KEGG [8] and GO [9].

**Network Construction and Analysis**

To unravel the system's behavior of genetic interactions and regulation of oxidative-stress responsive gene, we established the oxidative-stress specific protein–protein interaction (PPI) network fused with DEGs and the 1st neighbors to the largest publicly available Escherichia coli str. K-12 substr. MG1655 PPI network from the STRING database [10]. We had an oxidative-stress specific PPI network with 315 nodes with 5,796 edges with combined score > 800. Next, we extracted the latest gene regulatory network (GRN) through RegulonDB [11]. We focused our exploration on the genes with maximum variance in gene expression and their 1st neighbors. We constructed the GRN with 310 genes and TFs connected with 576 edges. The GRN has positive (+), negative (-), and positive or negative (+-; undetermined) regulatory direction from TF to a gene. To comprehensively understand the interactions and gene regulation, we merged the PPI and GRN with resulting in 444 nodes and 6,347 edges, including interactions and regulation [12]. The network centrality analyses were performed on the merged network to identify the most important regulators and players during oxidative stress. The resultant networks were visualized in Cytoscape [13].

**Supplemental Figures:**

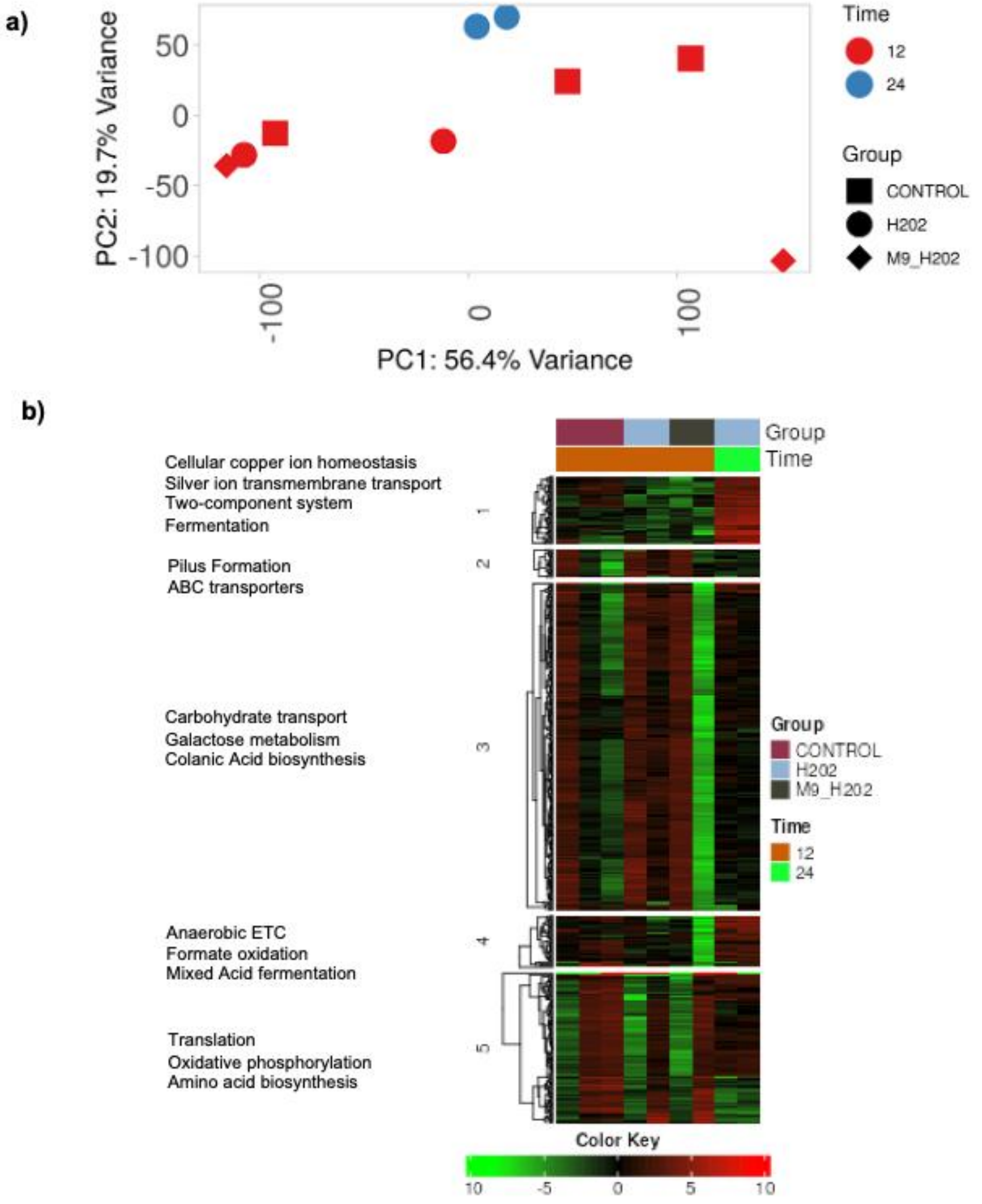


**Figure S1: The Transcriptome variability in control and $H_2O2$ treated E. coli**. a) The PCA plot covering ~75% of variance in the entire dataset. The control and $H_2O_2$ treated samples are well segregated. b) The gene expression heatmap of 500 most variable genes in the data. The clusters are determined by k-means, The functional enrichment was done by GO, KEGG, and EcoCys.

**References**

1 Reynolds, J. Serial Dilution Protocols. (American Society of Microbiology, 2005).
2 Hudzicki, J. Kirby Bauer Disk Diffusion Susceptibility Test Protocol. (American Society for Microbiology, 2009).
3 O'Toole, G. A. Microtiter dish biofilm formation assay. *J Vis Exp*, doi:10.3791/2437 (2011).
4 Arora, N. K. & Verma, M. Modified microplate method for rapid and efficient estimation of siderophore produced by bacteria. *3 Biotech* **7**, 381, doi:10.1007/s13205-017-1008-y (2017).
5 Zorraquino, V., Kim, M., Rai, N. & Tagkopoulos, I. The Genetic and Transcriptional Basis of Short and Long Term Adaptation across Multiple Stresses in Escherichia coli. *Mol Biol Evol* **34**, 707-717, doi:10.1093/molbev/msw269 (2017).
6 Ewels, P. A. *et al.* The nf-core framework for community-curated bioinformatics pipelines. *Nat Biotechnol* **38**, 276-278, doi:10.1038/s41587-020-0439-x (2020).
7 Love, M. I., Huber, W. & Anders, S. Moderated estimation of fold change and dispersion for RNA-seq data with DESeq2. *Genome Biology* **15**, doi:ARTN 550

10.1186/s13059-014-0550-8 (2014).
8 Kanehisa, M. & Goto, S. KEGG: kyoto encyclopedia of genes and genomes. *Nucleic Acids Res* **28**, 27-30, doi:10.1093/nar/28.1.27 (2000).
9 Ashburner, M. *et al.* Gene ontology: tool for the unification of biology. The Gene Ontology Consortium. *Nat Genet* **25**, 25-29, doi:10.1038/75556 (2000).
10 Szklarczyk, D. *et al.* STRING v10: protein-protein interaction networks, integrated over the tree of life. *Nucleic Acids Res* **43**, D447-452, doi:10.1093/nar/gku1003 (2015).
11 Salgado, H. *et al.* RegulonDB v12.0: a comprehensive resource of transcriptional regulation in E. coli K-12. *Nucleic Acids Res* **52**, D255-D264, doi:10.1093/nar/gkad1072 (2024).
12 Mishra, B., Sun, Y., Howton, T. C., Kumar, N. & Mukhtar, M. S. Dynamic modeling of transcriptional gene regulatory network uncovers distinct pathways during the onset of Arabidopsis leaf senescence. *NPJ Syst Biol Appl* **4**, 35, doi:10.1038/s41540-018-0071-2 (2018).
13 Shannon, P. *et al.* Cytoscape: a software environment for integrated models of biomolecular interaction networks. *Genome Res* **13**, 2498-2504, doi:10.1101/gr.1239303 (2003).
